# Subtle immunological differences in mRNA-1273 and BNT162b2 COVID-19 vaccine induced Fc-functional profiles

**DOI:** 10.1101/2021.08.31.458247

**Authors:** Paulina Kaplonek, Deniz Cizmeci, Stephanie Fischinger, Ai-ris Collier, Todd Suscovich, Caitlyn Linde, Thomas Broge, Colin Mann, Fatima Amanat, Diana Dayal, Justin Rhee, Michael de St. Aubin, Eric J. Nilles, Elon R. Musk, Anil S. Menon, Erica Ollmann Saphire, Florian Krammer, Douglas A. Lauffenburger, Dan H. Barouch, Galit Alter

## Abstract

The successful development of several COVID-19 vaccines has substantially reduced morbidity and mortality in regions of the world where the vaccines have been deployed. However, in the wake of the emergence of viral variants, able to evade vaccine induced neutralizing antibodies, real world vaccine efficacy has begun to show differences across the mRNA platforms, suggesting that subtle variation in immune responses induced by the BNT162b2 and mRNA1273 vaccines may provide differential protection. Given our emerging appreciation for the importance of additional antibody functions, beyond neutralization, here we profiled the post-boost binding and functional capacity of the humoral response induced by the BNT162b2 and mRNA-1273 in a cohort of hospital staff. Both vaccines induced robust humoral immune responses to WT SARS-CoV-2 and VOCs. However, differences emerged across epitope-specific responses, with higher RBD- and NTD-specific IgA, as well as functional antibodies (ADNP and ADNK) in mRNA-1273 vaccine recipients. Additionally, RBD-specific antibody depletion highlighted the different roles of non-RBD-specific antibody effector function induced across the mRNA vaccines, providing novel insights into potential differences in protective immunity generated across these vaccines in the setting of newly emerging VOCs.

## Introduction

The unprecedented rapid development of multiple SARS-CoV-2 vaccines marked a breakthrough in vaccine development and provided hope for an end to the COVID-19 pandemic. However, rising numbers of breakthrough infections, driven by evolving variants of concern in the setting of waning immunity, have clearly illustrated the urgent need to define correlates of immunity. Preliminary immune correlates analyses have shown a strong relationship between neutralizing antibody levels and vaccine efficacy ^1^. Yet, surprisingly, antibody binding titers provide even stronger surrogate of protection across vaccine platforms ^2–4^, with protection observed prior to the evolution of neutralizing antibodies ^4, 5^, and has persisted even in the setting of waning neutralizing antibodies ^6^. These data argue for a potential role for alternative protective antibody mechanisms of action.

Beyond their role in binding and neutralization, antibodies mediate a wide array of additional protective immunological functions through their ability to recruit the immune system via Fc receptors and complement ^7^. Fc-effector functions have been implicated in protection against multiple pathogens, including influenza ^8^, anthrax ^9^, HIV ^10^, malaria ^11^, and the Ebola virus ^12^. Likewise, Fc-effector functions have been linked to protection against SARS-CoV-2 both following vaccination and administration of monoclonal therapeutics in animal models ^13–15^. Importantly, Fc-effector function has been implicated in reducing the severity of disease rather than transmission, and thus likely to play a more critical role in vaccine attenuated disease, rather than simple blockade of infection. While accumulating data points to the ability of adenoviral platforms to evoke strong Fc-effector functions ^16, 17^ that have been linked to protection against HIV or malaria ^18^, less is known about the ability of newer vaccine platforms, including the mRNA vaccine, in eliciting these functions.

Striking protection was observed in the phase 3 BNT162b2 ^4^ and mRNA-1273 ^19^ studies, with 94.1% and 95% vaccine efficacy observed at a time when the D614G strain was in circulation. Yet, despite the similar antibody titers and neutralizing antibody levels induced by these mRNA vaccines, emerging real world data have begun to point to differences in the real world efficacy of these two vaccines. Specifically, in the face of the Delta-variant, ~40% and ~75% efficacy has been noted in BNT162b2 and mRNA-1273 vaccinees ^20^. Preliminary data in pregnant women have begun to point to differences in vaccine induced humoral immune responses elicited by the Pfizer/BioNTech (BNT162b2), and Moderna (mRNA-1273) vaccines ^21^ proposed to be driven by differences in vaccine dose, formulation, and/or the one week-delay in boosting ^22^. However, whether similar differences exist in the non-pregnant vaccinees, particularly across VOCs, remains incompletely understood.

Thus, here we compared the humoral response across the BNT162b2 and mRNA-1273 at the peak immunogenicity in a group of hospital staff. Both vaccines induced robust functional humoral immune responses, yet differences were noted in the vaccine induced antibody profiles across the vaccine groups, with higher RBD- and NTD-specific IgA, as well as functional antibodies (ADNP and ADNK) among mRNA-1273 immunized vaccinees. Both mRNA vaccines drove robust responses across the VOCs, able to bind to multiple FcRs, including the beta and delta variants. Moreover, RBD-specific antibody depletion highlighted the presence of non-RBD-specific antibody effector function deployed by both platforms, albeit at different levels, providing an explanation for differential sustained levels of Fc-effector function observed in the face of evolving VOCs.

## Methods

### Study populations

#### mRNA-vaccinated hospital staff

The Beth Israel Deaconess Medical Center institutional review board approved this study (#2021P000344) and the parent biorepository study (#2020P000361); participants provided written informed consent. We conducted a descriptive cohort study of hospital staff ≥ 18 years of age who were planning to receive an mRNA COVID-19 vaccine from December 2020 through February 2021 using samples collected in a larger hospital-wide, prospective data and tissue biorepository. Participants self-referred from flyers posted in the hospital vaccine clinics. All participants provided blood samples collected close to each vaccine dose and two to eight weeks after the second dose for the mRNA-1273 (Moderna) or BNT162b2 (Pfizer/BioNTech) COVID-19 vaccine. The analysis presented here includes non-pregnant individuals without immunosuppression medication use. To further characterize the study population, participants were asked to provide their race and ethnicity based on specified categories for each; they could select multiple race categories. Participants also reported if they had fever symptoms following either vaccine dose.

#### Community-acquired COVID-19 individuals

Industry employees (Space Exploration Technologies Corp.) were volunteer tested for COVID-19, starting in April 2020. Participants completed a study survey including the collection of COVID-19 related symptoms. The cohort largely included mild-symptomatic infections ^45^. Upon obtaining informed consent, blood samples were collected and used for immune profiling. The median age of the seropositive population was 32 years (range 19–62 years), and 84% were males. The enrolled participants were 66% White, 8% Asian, 6% More than one race, 2% Black, 1% American Indian/Alaska Native and 17% unknown. Volunteers were tested by PCR and for antibodies monthly. All antibody positive individuals were included in the study^45^.

Both studies were approved by the Massachusetts General Brigham Healthcare (previously Partners Healthcare) Institutional Review Board. All participants provided written informed consent.

### Antigens

Antigens used for Luminex based assays: SARS-CoV-2 D614G WT S (kindly provided by Erica Saphire, La Jolla Institute for Immunology), SARS-CoV-2 S1 (Sino Biological), SARS-CoV-2 S2 (Sino Biological) and SARS-CoV-2 RBD (kindly provided by Aaron Schmidt, Ragon Institute) as well as SARS-CoV-2 VOC,s such as Alpha B.1.1.7 S (LakePharma), Beta B.1.351 S (LakePharma), Gamma P1 S (LakePharma), Epsilon B.1.427 (kindly provided by Erica Saphire, La Jolla Institute for Immunology), Iota B. 1.526 (LakePharma), Kappa B. 1.617.1 S (Sino Biological) and Delta B.1.617.2 S (kindly provided by Erica Saphire, La Jolla Institute for Immunology) and Alpha B.1.1.7, Beta B.1.351, Gamma P1, Kappa B.1.617.2 and Delta B.1.617.2 RBDs (kindly provided by Florian Krammer, Icahn School of Medicine at Mount Sinai).

### Luminex profiling

Serum samples were analyzed by customized Luminex assay to quantify the relative concentration of antigen-specific antibody isotypes, subclasses, and Fcγ-receptor (FcγR) binding profiles, as previously described ^46, 47^. Briefly, SARS-CoV-2 antigens were used to profile specific humoral immune responses. Antigens were coupled to magnetic Luminex beads (Luminex Corp) by carbodiimide-NHS ester-coupling (Thermo Fisher). Antigen-coupled microspheres were washed and incubated with plasma samples at an appropriate sample dilution (1:500 for IgG1 and all low affinity Fcγ-receptors, and 1:100 for all other readouts) for 2 hours at 37°C in 384-well plates (Greiner Bio-One). The high affinity FcR was not tested due to its minimal role in tuning antibody effector function ^48^. Unbound antibodies were washed away, and antigen-bound antibodies were detected by using a PE-coupled detection antibody for each subclass and isotype (IgG1, IgG3, IgA1, and IgM; Southern Biotech), and Fcγ-receptors were fluorescently labeled with PE before addition to immune complexes (FcγR2a, FcγR3a; Duke Protein Production facility). After one hour of incubation, plates were washed, and flow cytometry was performed with an iQue (Intellicyt), and analysis was performed on IntelliCyt ForeCyt (v8.1). PE median fluorescent intensity (MFI) is reported as a readout for antigenspecific antibody titers.

### Antibody-dependent complement deposition (ADCD)

Antibody-dependent complement deposition (ADCD) was conducted as previously described ^49^. Briefly, SARS-CoV-2 antigens were coupled to magnetic Luminex beads (Luminex Corp) by carbodiimide-NHS ester-coupling (Thermo Fisher). Coupled beads were incubated for 2 hours at 37°C with serum samples (1:10 dilution) to form immune complexes and then washed to remove unbound immunoglobulins. In order to measure antibody-dependent deposition of C3, lyophilized guinea pig complement (Cedarlane) was diluted in gelatin veronal buffer with calcium and magnesium (GBV++) (Boston BioProducts) and added to immune complexes. Subsequently, C3 was detected with an anti-C3 fluorescein-conjugated goat IgG fraction detection antibody (Mpbio). The flow cytometry was performed with iQue (Intellicyt) and an SLab robot (PAA). ADCD was reported as the median of C3 deposition.

### Antibody-dependent cellular (ADCP) and neutrophil (ADNP) phagocytosis

Antibody-dependent cellular phagocytosis (ADCP) and antibody-dependent neutrophil phagocytosis (ADNP) were conducted according to the previously described protocols ^50, 51^. In detail, SARS-CoV-2 antigens were biotinylated using EDC (Thermo Fisher) and Sulfo-NHS-LCLC biotin (Thermo Fisher) and coupled to yellow-green (505/515) fluorescent Neutravidinconjugated beads (Thermo Fisher), respectively. To form immune complexes, antigen-coupled beads were incubated for 2 hours at 37°C with 1:100 diluted serum samples and then washed to remove unbound immunoglobulins. For ADCP, the immune complexes were incubated for 16-18 hours with THP-1 cells (1.25×10^5^ THP-1 cells/mL) and for ADNP for 1 hour with RBC-lyzed whole blood. Following the incubation, cells were fixed with 4% PFA. For ADNP, RBC-lyzed whole blood was washed, stained for CD66b+ (Biolegend) to identify neutrophils, and then fixed in 4% PFA. Flow cytometry was performed to identify the percentage of cells that had phagocytosed beads as well as the number of beads that had been phagocytosis (phagocytosis score = % positive cells × Median Fluorescent Intensity of positive cells/10000). The Flow cytometry was performed with 5 Laser LSR Fortessa Flow Cytometer and analysis was performed using FlowJo V10.7.1.

### Antibody-dependent NK cell degranulation

Antibody dependent NK cell degranulation as described previously (49). Briefly, SARS-CoV-2 antigens were coated to 96-well ELISA at the protein concentration of 2 g/ml, incubated at 37°C for 2hrs and blocked with 5% BSA at 4°C overnight. NK cells were isolated from whole blood from healthy donors (by negative selection using RosetteSep (STEMCELL) then separated using a ficoll gradient. NK cells were rested overnight in media supplemented with IL-15. Serum samples were diluted at 1:25. After blocking, samples were added to coated plates and immune complexes were formed for two hours at 37°C. After the two hours, NK cells were prepared (antiCD107a–phycoerythrin (PE) – Cy5 (BD), brefeldin A (10 μg/ml) (Sigma), and GolgiStop (BD)), and added to each well. for 5 hours at 37°C. The cells were stained for surface markers using anti-CD3 PacBlue (BD), anti-CD16 APC-Cy5 (BD), and anti-CD56 PE-Cy7 (BD) and permeabilized. The flow cytometry was performed. NK cells were gates as CD3-, CD16+, CD56+ cells and NK cell activity was determined as the percent of NK cells positive for CD107a and MIP-1b.

### RBD-specific antibody depletion from polyclonal serum samples

SARS-CoV-2 RBD-coated magnetic beads (ACROBiosystems) were prepared according to the manufacturer’s protocol and resuspended in ultrapure water at 1 mg/ml concentration. Beads were washed three times in phosphate-buffered saline (PBS) with 0.05% bovine serum albumin (BSA) by using a magnet. Serum samples were incubated with beads, at 3:1 beads:serum ratio, rotating overnight at 4°C. A magnet was used to separate beads with surfaces-bounded RBD-specific antibodies from the RBD-specific antibodies-depleted supernatant. A mock depletion (pre-depletion samples) was performed by adding 150 μl of PBS + 0.05% BSA and incubating rotating overnight at 4°C. Functional assays were performed with pre- and post-depletion samples.

### Statistics

Data analysis was performed using R version 4.0.2 (2020-06-22). Comparisons between vaccination arms were performed using Mann-Whitney U-test test followed by Benjamini Hochberg (BH) correction. Antigen responses (e.g., wild type to alpha) were compared using Wilcoxon-signed rank test followed by BH. Comparisons between RBD+ and RBD-samples were done using paired t-test.

Prior to any multivariate analysis, all data were normalized using z-scoring. Multivariate classification models were trained to discriminate between individuals vaccinated with BNT162b2 and individuals vaccinated with mRNA-1273 using all the measured antibody responses. Models were built using a combination of the least absolute shrinkage and selection operator (LASSO) for feature selection and then classification using partial least square discriminant analysis (PLS-DA) with the LASSO-selected features using R package “ropls” version 1.20.0 (Thévenot et al., 2015) and “glmnet” version 4.0.2. Model accuracy was assessed using ten-fold cross-validation. For each test fold, LASSO-based feature selection was performed on logistic regression using the training set for that fold. LASSO was repeated 100 times, features selected at least 90 times out of 100 were identified as selected features. PLS-DA classifier was applied to the training set using the selected features and prediction accuracy was recorded. Selected features were ordered according to their Variable Importance in Projection (VIP) score and the first two latent variables (LVs) of the PLS-DA model were used to visualize the samples. A co-correlate network analysis was carried out to identify features that highly correlate with the LASSO selected features, and thus are potentially equally important for discriminating the vaccination arms. Correlations for the co-correlate network were performed using Spearman method followed by Benjamini-Hochberg multiple correction ^52^. The co-correlate network was generated using R package “network” version 1.16.0 ^53^. All other figures were generated using ggplot2 ^54^.

## Results

### Enrollment

Seventy-three participants, 28 receiving mRNA-1273 and 45 receiving BNT162b2, were included from the hospital-wide biorepository of vaccinated individuals who received an mRNA COVID19 vaccine and had serum available for analysis following their second vaccine dose (Table 1). The vaccines were delivered intramuscularly, 30 μg of BNT162b2 and 100 μg of mRNA-127 were delivered, three and four weeks apart, respectively. Samples were obtained a median (interquartile range, IQR) of 19 (15, 26) days after the second vaccine dose. Prior SARS-CoV-2 infection (mild disease) was diagnosed in 7% of mRNA-1273 vaccinated and 2% of BNT162b2 vaccinated. After the second dose, fever was reported in 12 (48%) mRNA-1273 vaccinated and 19 (45%) of BNT162b2 vaccinated participants (Table 1).

**Table 1.** Characteristics of vaccinated participants.

|  | mRNA-1273<br>vaccinated<br>n = 28 | BNT162b2<br>vaccinated<br>n = 45 |
| --- | --- | --- |
| Age, median (IQR), yrs | 32 (26, 42) | 32 (27, 37) |
| Sex at birth, female | 23 (82) | 41 (91) |
| Race, No. | 25 | 42 |
| White | 20 (80) | 29 (69) |
| Black | 0 | 5 (12) |
| Asian | 2 (8) | 3 (7) |
| Multi-racial | 2 (8) | 4 (10) |
| Other | 2 (8) | 1 (2) |
| Ethnicity, No. | 26 | 41 |
| Hispanic or Latino | 3 (12) | 6 (15) |
| Chronic medical conditions |  |  |
| Diabetes | 0 | 1 (2) |
| Hypertension | 2 (7) | 3 (7) |
| Obesity (body mass index $\geq 30\text{mg/kg}^2$) | 3 (11) | 2 (3) |
| Asthma | 3 (11) | 3 (7) |
| Prior SARS-CoV-2 infection | 2 (7) | 1 (2) |
| Days from second vaccine dose to sample collection, median (IQR) | 17 (15-20) | 21 (17-27) |
| Fever within 48 hours (by self-report) |  |  |
| After first dose, n/N (%) | 1/24 (4) | 0/41 (0) |
| After second dose, n/N (%) | 12/25 (48) | 19/41 (46) |

IQR: interquartile range
Participants were chosen from the biorepository based on their sample availability at the time of the analysis.

### mRNA-1273 and BNT162b2 COVID-19 vaccines induce robust WT-antibody Fc-profiles

The two approved mRNA vaccines clearly induce robust antibody titers and neutralization ^23, 24^, however, real world efficacy data has begun to show differences across the vaccines in their ability to resist delta viral infection ^20^. Specifically, approximately 42% protection has been noted following the BNT162b2 vaccine, and 76% efficacy has been observed with the mRNA-1273 vaccine. Recent profiling in pregnant women highlighted significant differences in the Fc-quality of the humoral immune responses induced by these two vaccines ^25^, potentially related to differences in dosage, formulation, the timing of the boost, or mRNA design. However, whether these same immunological changes occur in non-pregnant vaccinees and vary across emerging variants of concern (VOCs) remains incompletely understood. Thus, here we sought to determine whether the two authorized COVID-19 mRNA vaccines elicit similar Fc profiles in a cohort of 73 health care workers vaccinated with either of the vaccines. The vaccines were delivered intramuscularly, 30 μg of BNT162b2 and 100 μg of mRNA-127 were delivered, three and four weeks apart, respectively, and blood was collected two weeks post the final immunization. The wild-type SARS-CoV-2 RBD-, N-terminal domain (NTD)-, S-, S1- and S2-specific antibody titers, Fc-receptor binding, and Fc-functions were analyzed.

Robust vaccine induced antibody responses were observed across both the mRNA-1273 (*n = 28*) and BNT162b2 (*n = 45*) vaccines (Figure. 1A), marked by slightly higher responses in the mRNA-1273 vaccinees. Univariate comparisons across each antigen and Fc-profile measurement highlighted the presence of equivalent IgG and IgM binding titers, but higher levels of IgA-binding titers elicited by the mRNA-1273 vaccine, particularly to the Spike, RBD, NTD, and S1 domains (Figure. 1B). Moreover, robust and largely equivalent cross Fc-receptor binding was observed across both vaccines, with the exception of enhanced NTD-specific Fc-receptor binding antibodies induced by the mRNA-1273 vaccine. Similarly, equivalent levels of antibody dependent complement deposition (ADCD) and antibody dependent cellular phagocytosis by monocytes (ADCP) were observed across the two vaccine groups at peak immunogenicity. Conversely, mRNA-1273 vaccinated individuals exhibited significantly higher levels of antibody dependent neutrophil phagocytosis (ADNP) and antibody dependent NK cell activation (degranulation: CD107a, cytokine secretion: IFND□, and chemokine secretion: MIP1□) (Figure. 1C).

**Figure. 1.**
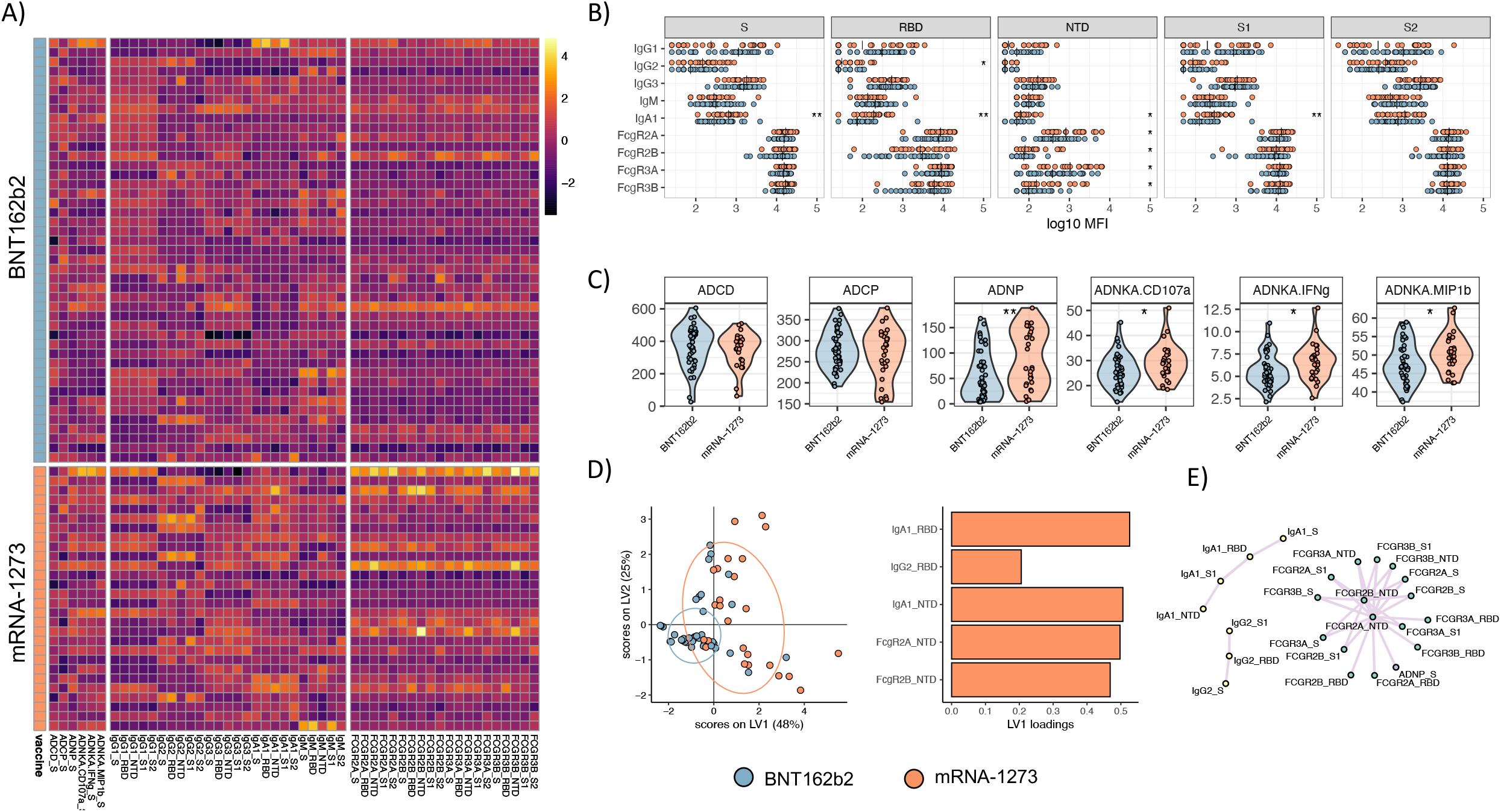
mRNA-1273 and BNT162b2 COVID-19 vaccines induce similar SARS-CoV-2 D614G WT antibody profiles. **(A)** Heatmap summarizes SARS-Cov-2 WT S, RBD, NTD, S1, and S2-specific IgG1, IgG2, IgG3, IgA1, and IgM titers, Fc measurements, and functional assays such, Antibody-dependent complement deposition (ADCD), cellular phagocytosis (ADCP) and neutrophil phagocytosis (ADNP), and NK cell activation (ADNKA, CD107a surface expression, and MIP-1β or IFNγ production) for each subject in BNT162b2 or mRNA-1273 arm. Titers and FcRs were first logtransformed and all measurements were z-scored. **(B-C)** Univariate comparisons of **(B)** antibody titers and FcRs binding for SARS-CoV-2 WT S, RBD, NTD, S1 and S2, as well as **(C)** S-specific antibody-mediated effector functions, such as ADCD, ADCP, ADNP and ADNKA, between BNT162b2 (blue) and mRNA-1273 (red) vaccines recipients. Mann-Whitney U-test corrected for multiple comparisons by the Benjamini-Hochberg (BH) method, was used. The adjusted *p* < 0.001 ***, *p* < 0.01 **, *p* < 0.05 *. **(D)** The least Absolute Shrinkage Selection Operator (LASSO) was used to select antibody features that contributed most to the discriminate subjects vaccinated with BNT162b2 or mRNA-1273. A partial least square discriminant analysis (PLSDA) was used to visualize samples. LASSO selected features were ranked based on their Variable of Importance (VIP) score, and the loadings of the latent variable 1 (LV1) were visualized in a bar graph. **(E)** A co-correlate network of the LASSO selected features was built using a threshold of absolute Spearman rho greater than 0.7 and BH-adjusted p-value lower than 0.01. Nodes were colored based on the type of measurement; titers, FcRs, and functional measurements. All shown links are positive correlations.

Given the univariate differences, we next aimed to define whether differences existed in the overall multivariate vaccine profile across the two vaccine groups. Thus, we used the least absolute shrinkage and selection operator (LASSO) feature selection to initially reduce all antibody features to a minimal set which represented the overall variation in the antibody profiles and to avoid over-fitting, followed by classification using partial least squares discriminant analysis (PLS-DA). Separation was observed across the two different mRNA vaccine profiles (Figure. 1D), marked largely by augmented responses in the mRNA-1273 vaccine induced immune response. Specifically, five features were selectively enhanced in the mRNA-1273 vaccine profiles, including RBD-specific IgA1 and IgG2, as well as NTD-specific IgA1, FcγR2A, and FcγR2B. Given the highly correlated nature of the vaccine induced humoral immune response, a correlation network analysis was built between LASSO-selected features and the overall immune response to define the additional features that may shift differentially across the vaccine profiles (Figure. 1E). Three clusters appeared, bearing elevated IgA response across all antigenic determinants, a small network of IgG2 responses, and a large network of Fc-receptor binding antibody responses across multiple antigenic targets all enriched among mRNA-1273 immunized individuals. These data point to robust humoral immune responses induced by both mRNA platforms, but that do diverge with enhanced epitope spreading, IgA immunity, and specific antibody effector functions in mRNA-1273 immunized individuals.

### mRNA-1273 and BNT162b2 vaccines induce FcR-binding responses to multiple VOCs

Despite the remarkable efficacy of the mRNA vaccines against the original SARS-CoV-2 variant, waves of variants have emerged that include amino acid substitutions that significantly diminish neutralizing antibody activity ^26–28^. Among the variants of concern (VOCs), the mRNA vaccines appear to neutralize Alpha (B.1.1.7) and Gamma (P.1) with only a minimal loss of activity, but exhibit compromised neutralizing activity against the beta (B.1.351) variant ^30 31^. Yet, whether Fc responses were equally affected across the VOCs remains unclear. Both mRNA-1273 and BNT162b2 vaccine-induced antibodies bound equally well across the Alpha B.1.1.7, Beta B.1.351, and Gamma P.1 VOCs (Figure. 2A). Interestingly, IgM titers were higher in BNT162b2 vaccinated individuals to the Beta and Gamma VOCs. Additionally, a trend towards higher IgG1-binding titers was also noted in the BNT162b2 immunized individuals across all 3 VOCs. Conversely, IgA responses were amplified in the mRNA-1273 immunized individuals (Figure. 2A). However, Fc receptor-binding antibodies were induced by both vaccines to all 3 VOCs at equivalent levels. Along the same lines, both antibody-dependent monocyte (ADCP) and neutrophil (ADNP) phagocytosis were largely equivalent across the variants (Figure. 2B), highlighting the robust Fc binding and functional profiles across VOCs elicited by both mRNA platforms. Yet, despite these univariate results, we finally aimed to ask if any multivariate differences could be observed across the two mRNA platforms in their VOC response (Figure. 2C). The LASSO/PLSDA revealed separation in the Fc-profiles induced to the VOCs between the mRNA-1273 and BNT162b vaccinated individuals (Figure. 2C). The profile was marked by higher IgM-Beta levels in BNT162b vaccinated individuals. Conversely, higher levels of IgA/IgG2 responses were observed to alpha in mRNA-1273 vaccinated individuals who also had higher levels of FCR2A and FCR2B NTD-binding antibodies. The extended LASSO co-correlate network further highlighted the presence of IgG2, IgA, and IgM-only networks across multiple VOCs, suggesting that isotype biased selection across the mRNA platform includes reactivities across VOCs. Additionally, a large network of highly functional pan VOC and epitope responses were observed in the mRNA1273 profile, marked by an enrichment of NTD-specific antibody responses. Thus, the two mRNA platforms elicit overall similar levels of functional antibodies to the VOCs, with an IgM/IgG biased profile induced by BNT162b2 and a more class switched IgA/IgG-driven profile induced by mRNA-1273.

**Figure. 2.**
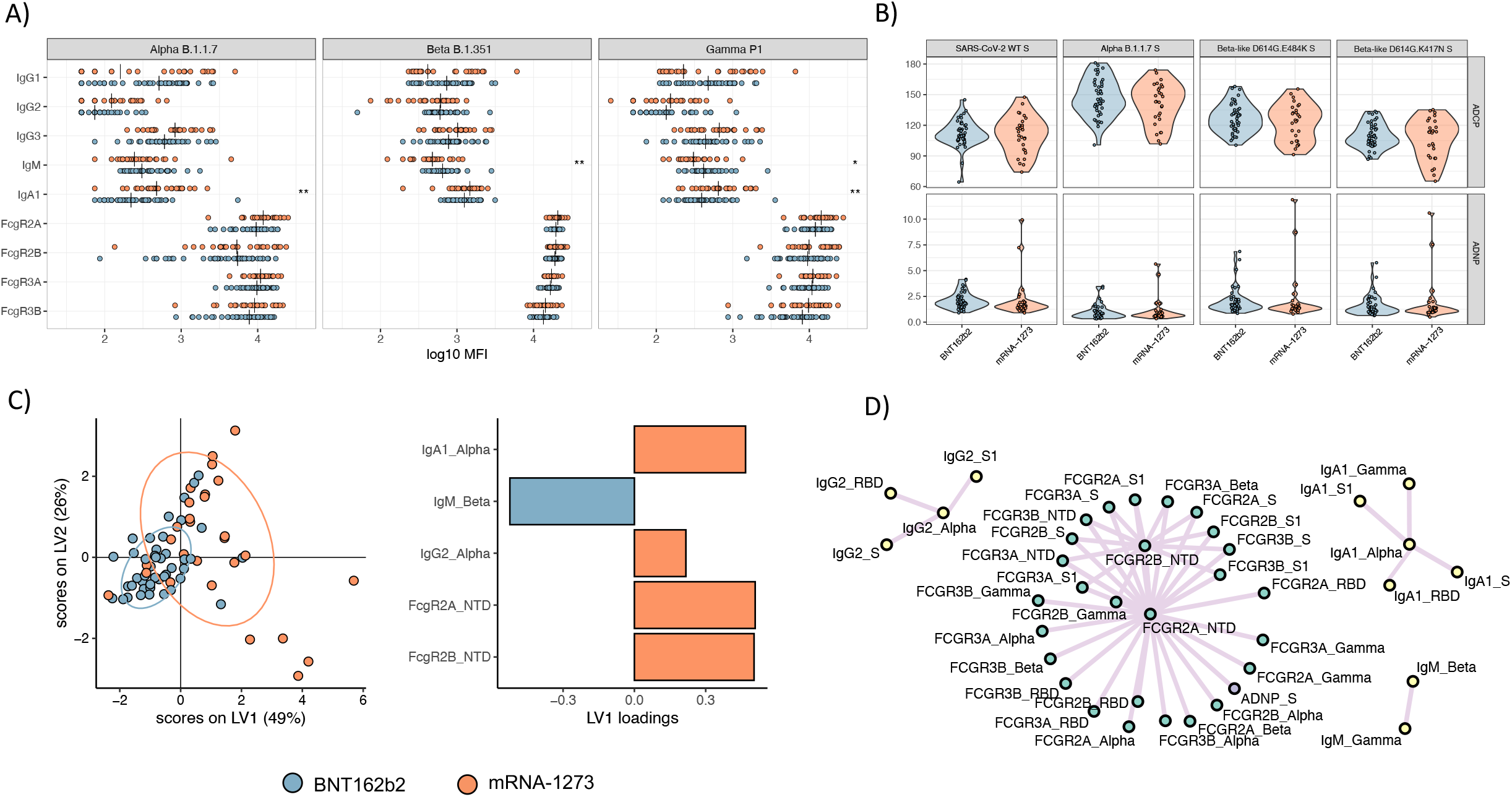
mRNA-1273 and BNT162b2 vaccines induce a comparable antibody profile across alpha, beta, and gamma SARS-CoV-2 VOCs. **(A-B)** Univariate comparisons of antibody titers and FcRs binding for SARS-CoV-2 VOCs, such as Alpha B.1.1.17, Beta B.1.351 and Gamma P1, as well as **(B)** antibody-dependent, cellular (ADCP) and neutrophil phagocytosis (ADNP), in BNT162b2 (blue) and mRNA-1273 (red) vaccinated individuals. Mann-Whitney U-test corrected for multiple comparison with the Benjamini-Hochberg (BH) method was used. The adjusted *p* < 0.001 ***, *p* < 0.01 **, *p* < 0.05 *. **(C)** LASSO-PLSDA model including all variant measurements. LASSO selected features were ranked based on their Variable of Importance (VIP) score, and the loadings of the latent variable 1 (LV1) were visualized in a bar graph. **(D)** A co-correlate network of the LASSO selected features was built using a threshold of absolute Spearman rho greater than 0.7 and BH-adjusted p-value lower than 0.01. Nodes were colored based on the type of measurement; titers, FcRs, and functional measurements. All shown links are positive correlations.

### mRNA-1273 and BNT162b2 vaccines induce more robust RBD and full Spike VOC-targeting Fc-functional antibodies compared to natural infection

Alarmingly, several variants, including the Alpha (B.1.1.7), Beta (B.1.351), Gamma (P1), and Delta (B.1.617.2) VOCs, have begun to breakthrough natural ^32^ and vaccine induced immune responses ^26^, causing large numbers of outbreaks. The unexpected significant reduction of effectiveness of mRNA vaccines against VOCs, especially delta (B.1617.2) variant, is emerging ^20^, albeit with the majority of breakthroughs remaining largely non-lethal ^1^. Yet, differential real-world efficacy against the Delta VOC ^20^ points to a nuanced immune response to delta. Thus, we next aimed to compare the cross-VOC antibody Fc-profiles targeting both VOC RBDs or full Spike antigens across a subset of the vaccinees and a group of mild-community acquired convalescent individuals. Antibody profiles were compared across mRNA-1273 (*n = 16*) and BNT162b (*n = 15*) vaccines and 10 convalescents (Figure. 3). mRNA vaccine induced IgG1 responses were higher than convalescent responses to the original variant RBD (WT), Alpha (B.1.1.7), Beta (B.1.351), Kappa (B.1.617.1), and Delta (B.1.617.2), but were lower for the gamma (P.1) RBD. Similar patterns were observed across all Fc-receptor RBD-binding antibodies induced by the mRNA vaccines, that all showed superior binding to FcRs compared to convalescent antibodies that bound poorly to all RBD VOCs compared to the WT variant (Figure. 3A). Interestingly, slightly higher antibody binding was noted across RBD VOCs for mRNA-1273 immunized individuals compared to BNT162b2, albeit the pattern of recognition was the same. Conversely, IgG1 and IgG3 binding patterns to S-specific antibodies revealed enhanced mRNA-spike specific binding to nearly all VOC-full Spike antigens, except the kappa variant, compared to WT Spike-specific antibody binding (Figure. 3B). Importantly, all Spike-specific binding IgG responses were lower in convalescents compared to mRNA vaccinees, with slightly higher IgG1 and IgG3 binding noted to nearly all VOC Spikes. In contrast, Fc-receptor binding antibodies exhibited equivalent VOC-recognition across both mRNA vaccine platforms, highlighting potential functional differences in RBD- and Spike-specific antibodies induced across the platforms. Thus, despite the more variable FcR-binding profiles to RBD VOCs, stable FcRbinding was noted to most full spike VOCs. Given the persistent protection against delta in newly/recently vaccinated individuals ^1^, but enhanced breakthrough over time, these data may suggest that the presence of broad non-RBD-specific functional immunity may be key to protection, given that VOC-RBDs clearly are able to exploit RBD-specific Fc-vulnerabilities even across vaccine platforms.

**Figure. 3.**
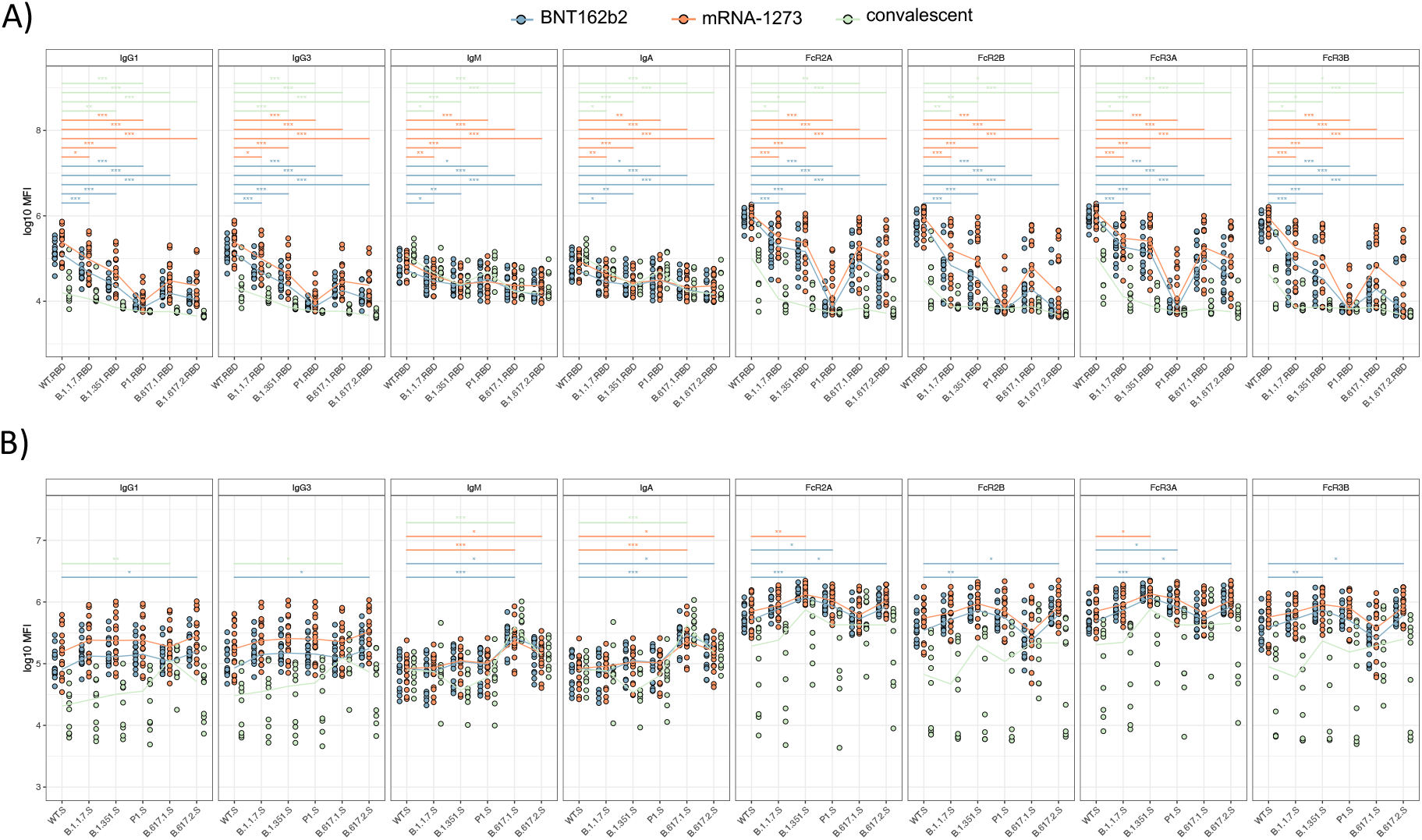
mRNA vaccines are able to induce the high level of Spike-specific antibody binding to FcRs across all SARS-CoV-2 VOCs. **(A-C)** Univariate comparisons of **(A)** RBD-specific, **(B)** S-specific IgG1, IgG2, IgG3, IgA1, and IgM titers and FcR bindings across SARS-CoV-2 WT and VOCs, such as Alpha B.1.1.7, beta B.1.351, Gamma P1, Kappa B.1.617.1 and Delta B.1.617.2, between the BNT162b2 (blue) and mRNA-1273 (red) vaccine recipients and convalescent individuals (green). **(C)** Delta B.1.617.2 S- and RBD-specific antibody titers and FcR binding between BNT162b2 (blue) and mRNA-1273 (red) vaccinees and convalescent individuals (green). Comparisons were made for each variant using Wilcoxon rank-sum test and corrected for multiple comparison using the BenjaminiHochberg (BH) method. The adjusted *p* < 0.001 ***, *p* < 0.01 **, *p* < 0.05 *.

### RBD-specific antibody depletion influences antibody-mediated monocyte and neutrophil phagocytosis

The data above suggested that potential differences in RBD and Spike specific contributions to polyclonal antibody Fc-binding profiles and function. Thus, to address this possibility, RBD-specific antibodies were depleted from the polyclonal serum of our vaccinees and convalescent samples (Figure. 4) and tested for opsonophagocytic functions linked to the natural resolution of infection ^33, 34^. Specifically, monocyte phagocytosis (ADCP, Figure. 4A) or neutrophil phagocytosis (ADNP, Figure. 4B) were evaluated across VOCs. RBD depletion resulted in a significant loss of ADCP against the WT variant Spike (Figure. 4A, bottom left). However, RBD depletion did not affect the ADCP response in convalescents to all other Spike VOCs, arguing that antibodies in natural infection that drive function largely target areas outside of the RBD. Conversely, RBD depletion resulted in slightly reduced ADCP in BNT162b2 immunized individuals to the WT, beta, and epsilon variants, but not to the alpha and gamma variants. Similarly, RBD-depletion resulted in reduced Beta and Gamma ADCP in mRNA-1273 immunized individuals. These data argue for variable, but low-level alterations in RBD-specific ADCP activity across VOC Spikes following natural infection, and differentially following mRNA vaccination, suggesting that non-RBD-specific antibodies may continue to drive opsonophagocytic control of the virus even in the setting of profound changes in the RBD that may knock out neutralization.

**Figure. 4.**
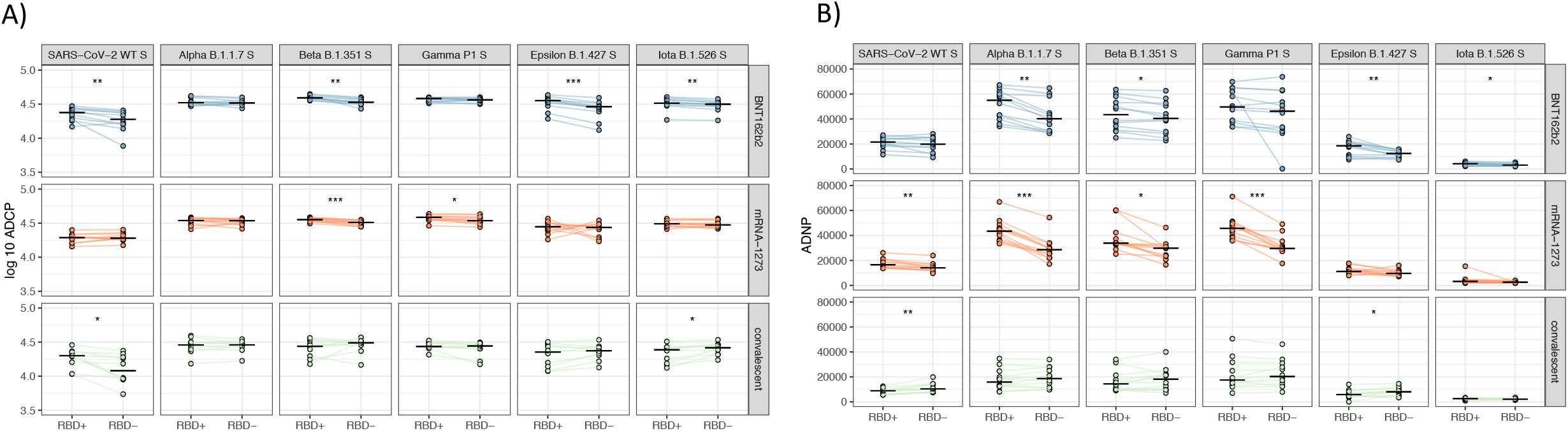
RBD-specific antibody depletion differently influences the antibody-mediated monocytes and neutrophils phagocytosis across SARS-CoV-2 VOCs. **(A)** Comparisons of antibody-mediated cellular (ADCP) and neutrophil (ADNP) phagocytosis across SARS-CoV-2 WT and VOC, such as Alpha B.1.1.7, beta B.1.351, Gamma P1, Epsilon B.1.427 and Iota B.1426 Spikes, between pre-depletion (RBD+) and post RBD-specific antibody depletion (RBD-) serum samples across from BNT162b2 (blue) and mRNA-1273 (red) vaccine recipients and convalescent individuals (green). The paired t-test corrected for multiple comparison by the Benjamini-Hochberg (BH) method was used. The adjusted *p* < 0.001 ***, *p* < 0.01 **, *p* < 0.05 *.

In contrast to ADCP, more variation was observed in neutrophil phagocytosis (ADNP) with RBD depletion. Interestingly, RBD depletion led to increased ADNP against the WT and Epsilon variants, suggesting that RBD-specific antibodies may, in fact, block this activity following the natural resolution of disease. Conversely, RBD depletion led to reduced ADNP to the alpha, beta, epsilon, and iota variants in BNT162b2 immunized individuals. Similarly, RBD depletion reduced ADNP to WT, alpha, beta, gamma, and iota in mRNA-1273 vaccinated individuals. Interestingly, the RBD depletion led to a more profound loss of ADNP in mRNA1273 immunized individuals compared to BNT162b2 immunized individuals, marking significant differences in the functional activity of particular sub-populations of antibodies elicited by these vaccines. Thus, given the higher levels of mRNA-1273 induced pan-VOC RBD specific immunity, that appears to contribute largely to ADNP activity, these data point to the possible existence of different epitope-specific functional correlates of immunity elicited across the two vaccine platforms and natural infection.

Thus, these data collectively show the robust induction of functional antibody responses that differ in their Fc-biology, following mRNA-1273 and BNT162b2 vaccination, marked by differences in the overall isotype/subclass, Fc-receptor binding profiles, and epitope specific functions across VOCs, providing some potential explanation for differences in persistent protection afforded against by this newly emerging vaccine platform.

## Discussion

Despite the remarkable protective immunity observed in the BNT162b2 and mRNA-1273 vaccine phase 3 studies against the original SARS-CoV-2 variant, breakthrough infections are on the rise globally among vaccinees ^35^. Yet, although the rise of breakthrough infections, severe disease, hospitalization, and death remain low in most populations, fatality rates have begun to emerge in the elderly, prompting discussions on additional vaccine boost ^36^. While both mRNA COVID-19 vaccines induced comparable and robust antibody titers and neutralization, emerging data point to variable levels of real-world vaccine efficacy across the platforms potentially linked to differences in the formulation, design, boosting intervals, and dose ^20^. Thus, while both vaccines induced robust antibody titers and neutralization, understanding immunological differences across the vaccines may provide critical insights on immune correlates of protection to guide next generation vaccine design and to guide boosting. Among the proposed nonneutralizing antibody immune mechanisms of protection, T cells have been proposed as a critical arm in the control of SARS-CoV-2 due to their known role in the battle against many viruses ^37, 38^. Yet, additional mechanisms, such as the role of antibody-mediated effector function, have also been shown to play a critical role in vaccine-mediated protection against a broad array of pathogens ^7^, including SARS-CoV-2 ^39^. Thus, here we deeply probed the functional humoral immune response induced by distinct mRNA vaccine platforms and probed their Fc-functional performance across VOCs, demonstrating robust Fc-functional responses, albeit distinct, induced by both the BNT162b2 and mRNA-1273 vaccines, that elicit Fc-effector functions against most VOCs, including the beta and delta variants, despite the documented loss of neutralizing activity. These data point to a potential role for vaccine induced Fc-effector function in mRNA vaccine induced protection against disease mediated by VOCs, but also to nuances in the functional response to VOCs that may contribute to real world efficacy differences across the platforms.

While no difference in neutralizing activity has been reported across the BNT162b2 and mRNA1273 vaccines ^24^, some differences in isotype/subclass and Fc-functions were noted across the two platforms. Consistent with previous observations in pregnant women ^25^, elevated levels of IgA were noted following mRNA-1273, accompanied by higher levels of ADNP and NK cell activation. Conversely, an IgM/IgG bias was noted in BNT162b2 vaccinated individuals to VOCs, pointing to differences in class switching across the mRNA platforms. Whether these differences contribute to different efficacy, particularly over time as the response wanes, remains unclear but will be addressed in long-term follow-up breakthrough studies. Moreover, whether these differences are related to differences in lipid nanoparticle composition, mRNA dose, and delay in boosting remains unclear, but highlights the potential of mRNA vaccines to drive “tunable” Fc effector function, that may be selectively shaped to achieve enhanced and selective control over particular target pathogens and non-infectious diseases in the future.

The majority of mutations in the VOCs occur in N-terminal and RBD domains ^40^, which play a critical role in enhancing binding to ACE2. Given that neutralizing antibodies target these same sites, aimed at occluding ACE2-access, or aimed at compromising RBD-structure to prevent ACE2-interactions, these same mutations inadvertently compromise neutralizing antibody activity. Conversely, Fc-functional antibodies can target the whole surface of the Spike antigen, and thus are not compromised in the same manner as neutralizing antibodies by individual or clusters of mutations found to alter ACE2-binding. In fact, while a large part of neutralizing antibodies targets the RBD, the RBD depletion did not knock out Fc-effector function in convalescent individuals or BNT162b2 and mRNA-1273. Instead, vaccine induced RBD-specific antibodies contributed minimally to ADCP mediated activity across the VOCs, however RBD-specific antibodies contributed more to ADNP activity across the VOCs, suggesting that epitope-specific functional programming likely occurs across the mRNA vaccines, with a more prominent focus of ADNP on the RBD in mRNA-1273 immunized individuals. Moreover, depletion of RBD-specific antibodies from convalescent plasma resulted in improved ADNP to some variant Spike antigens, suggesting that RBD-specific antibodies may block Fc-effector function in natural infection, either by blocking access of additional functional antibodies, or due to altered or perturbed Fc glycosylation induced in natural infection. Thus, these data point to the vaccine and infection-induced differences in Fc-programming at an epitope-specific level, which may play a critical part in the level of protection against disease severity across naturally immune and vaccinated populations that have been observed across breakthrough cases globally. Thus, understanding the relationship between non-neutralizing antibody functions, epitope specificity, and clinical protection may provide unexpected insights to plan for rational boosting efforts to stop the continuous emergence of VOC.

The spread of the delta VOC has raised concerns globally about vaccine efficacy and the need for boosting. While hospitalizations remain consistently high in regions of the globe where vaccine deployment has been slow, most infections in previously vaccinated individuals do not require hospitalization. These data suggest that while both naturally and vaccine induced antibodies gradually lose the capability of preventing transmission against VOCs, vaccine antibodies may still provide some barrier to severity of disease. The data here demonstrate the ability of the BNT162b2 and mRNA-1273 vaccines to induce robust Fc-effector functions, previously linked to the resolution of severe disease in unvaccinated individuals ^34, 41, 42^, across many VOCs, despite the reported loss of neutralization. Thus, despite increased breakthroughs in BNT162b2, the quality of the recall, an anamnestic response may be sufficient to respond to SARS-CoV-2 and drive control and clearance of infection. However, understanding the differences in transmission blocking activity across the BNT162b2 and mRNA-1273 may provide new clues for the redesign of vaccines able to provide a longer barrier of protection against the virus, which may be required to slow the rates of evolution of variants of concern. Linked to our emerging appreciation for the role of Fc-effector function in protection from infection/disease in non-human primates ^13, 43^, hamsters ^14^, and mice ^44^, upcoming immune correlates analyses and breakthrough studies will provide a concrete recognition for the role of Fc-effector function in protection against SARS-CoV-2. Yet, ultimately, the data here point to the potential nuanced differences in the quality of the humoral immune response induced across the two authorized mRNA vaccine technologies, able to broadly harness multiple Fc-effector functions, and tune these functions differentially to distinct epitopes and innate immune cell types depending on dose, formulation, or booster-timing.

## Contributions

P.K., D.C., D.A.L., and G.A. analyzed and interpreted the data. P.K., S.F., C.L., T.B., and T.S. performed experiments. D.C. performed the analysis. A.Y.C. and D.H.B. managed and collected samples and managed the clinical data for vaccinated participants. D.D., J.R., A.S.M., E.J.N. and E.R.M. managed samples and data collection for the Community-acquired COVID-19 cohort. E.O.S. and F.K. produced SARS-CoV-2 antigens. G.A. supervised the project. P.K. and G.A. drafted the manuscript. All authors critically reviewed the manuscript.

